# Influence of visual feedback persistence on visuo-motor skill improvement

**DOI:** 10.1101/2021.01.26.428288

**Authors:** Alyssa Unell, Zachary M. Eisenstat, Ainsley Braun, Abhinav Gandhi, Sharon Gilad-Gutnick, Shlomit Ben-Ami, Pawan Sinha

## Abstract

Towards the larger goal of understanding factors relevant for improving visuo-motor control, we investigated the role of visual feedback for modulating the effectiveness of a simple hand-eye training protocol. The regimen comprised a series of curve tracing tasks undertaken over a period of one week by neurologically healthy individuals with their non-dominant hands. Our three subject groups differed in the training they experienced: those who received ‘Persistent’ visual-feedback by seeing their hand and trace evolve in real-time superimposed upon the reference patterns, those who received ‘Non-Persistent’ visual-feedback seeing their hand movement but not the emerging trace, and a ‘Control’ group that underwent no training. Improvements in performance were evaluated along two dimensions – accuracy and steadiness, to assess visuo-motor and motor skills, respectively. We found that persistent feedback leads to a significantly greater improvement in accuracy than non-persistent feedback. Steadiness, on the other hand, benefits from training irrespective of the persistence of feedback. Our results not only demonstrate the feasibility of rapid visuo-motor learning in adulthood, but more specifically, the influence of visual veridicality and a critical role for dynamically emergent visual information.

## Introduction

Our capacity to perform visually guided motor actions, also known as visuo-motor coordination (1), plays an important role in our ability to effectively interact with our surroundings. These skills typically undergo rapid improvement during early childhood. Here, we focus on the extent and mechanism of enhancing this skill in adulthood, an issue that is interesting from both a scientific and applied perspective.

One of the most extensively studied factors for modulating visuo-motor development and learning is visual-feedback as a source of online guidance for motor execution and planning (2–4). Human developmental progression exhibits a steady increase in the importance of visual feedback for motor control. In early childhood, when intentional reaching first emerges around 4 months of age, infants’ reaches are inefficient: The hand speeds up and slows down multiple times as it takes a circuitous route to the target. Although initially believed to result from over-correcting the hand’s trajectory based on visual feedback (5), more recent evidence suggests that the jerkiness of infants’ hand movements is not significantly modulated by visual feedback about the hand relative to the target. Infants reach to targets with or without sight of the hand (as when presented with a glowing target in a dark room) at the same age (6), and reaching kinematics are similar in conditions that permit or deny visual feedback (7,8). Past the neonatal stage, as infants engage in increasingly higher precision tasks, the use of visual feedback for guiding goal directed movement emerges rapidly (9,10). By around 15 months, reaches in the dark are less straight and take longer to complete compared to reaches in light, and like adults (11), children reach more slowly without visual feedback (7,9). A similar result of slower hand movement was found in congenitally blind individuals, while functional characteristics (coordination and kinematic profile) of their basic reach to grasp were preserved (11). Thus, the visuo-motor system develops to utilize real-time sensory feedback for making on-line assessments, allowing it to organize visuo-motor plans and improve movement efficiency (12).

Our focus here is on characterizing aspects of visual feedback during training that induce visuo-motor skill improvement. Past studies have explored the influence of visual feedback *presence* (manipulating visibility of the moving hand or its start and end position) and *timing* (feedback given during the movement or after its completion) on reaching, pattern reproduction and adaptation (13–18). A factor whose role remains largely unknown is the *persistence* of visual feedback. Two very different kinds of information are provided by persistent visual feedback of the movement trajectory: the instantaneous position information of the end effector, and information about earlier locations. Comparison of the current position with earlier positions yields dynamically emergent information about the ongoing movement sequence. The addition of a real time trace is therefore more significant than information about instantaneous position alone in providing useful visual information about movement kinematics (19,20).

We investigate pattern production through curve tracing, a type of movement in which persistent visual feedback naturally occurs in the form of the produced patterns (19,21–23). The specific question we pose is whether movement in response to reference patterns is enough to improve eye-hand coordination in fine motor activities, or if persistent feedback essential to this process.

We operationalize our investigation of visuo-motor skill improvement in the context of pattern tracing using an individual’s non-dominant hand. The non-dominant hand is typically not used for performing fine motor tasks like writing or drawing. This lack of natural practice in the non-dominant hand provides us an opportunity to examine visuo-motor improvements in healthy adults. In right-handed subjects, use of the left hand offers an additional advantage in enhancing visuo-motor skill learning, as the contralateral right-hemisphere has been described as dominating spatial processing, including aspects of visuomotor integration (24–26). The low level of baseline proficiency with the left hand not only allows latitude for improvement, but also eliminates the unintentional training that might occur during the training period if it were the dominant hand being tested, since it would be impractical to prevent usage over the course of multiple days.

We examine the improvement of right-handed individuals’ left-hand figure tracing proficiency along the dimensions of accuracy (how well does a tracing match a reference pattern?) and steadiness (how unwavering is the stroke making?). Accuracy is regarded as an expression of visuo-motor coordination and steadiness as an expression of general motor skill (27–32).

## Methods

Our experimental design involved having participants practice tracing over figures with or without persistent visual feedback. We compared their accuracy and steadiness metrics pre- and post-training. The study was approved by the MIT’s institutional review board, the Committee on the Use of Humans as Experimental Subjects, informed consent was obtained from all participants, and all experiments were performed in accordance with relevant guidelines and regulations.

### Stimuli for tracing

We used a collection of 330 letters drawn from several non-English alphabets, chosen to span a wide range of letter appearances (curved and straight strokes, varying degrees of letter complexity), and to be unfamiliar to the average participant in the US. A sample set of our stimuli are shown in Figure 1a. All letters were printed in gray, with four letters on each A4 page, such that on average, each letter spanned 10cm × 10cm, with a stroke width of 2mm.

**Figure 1:**
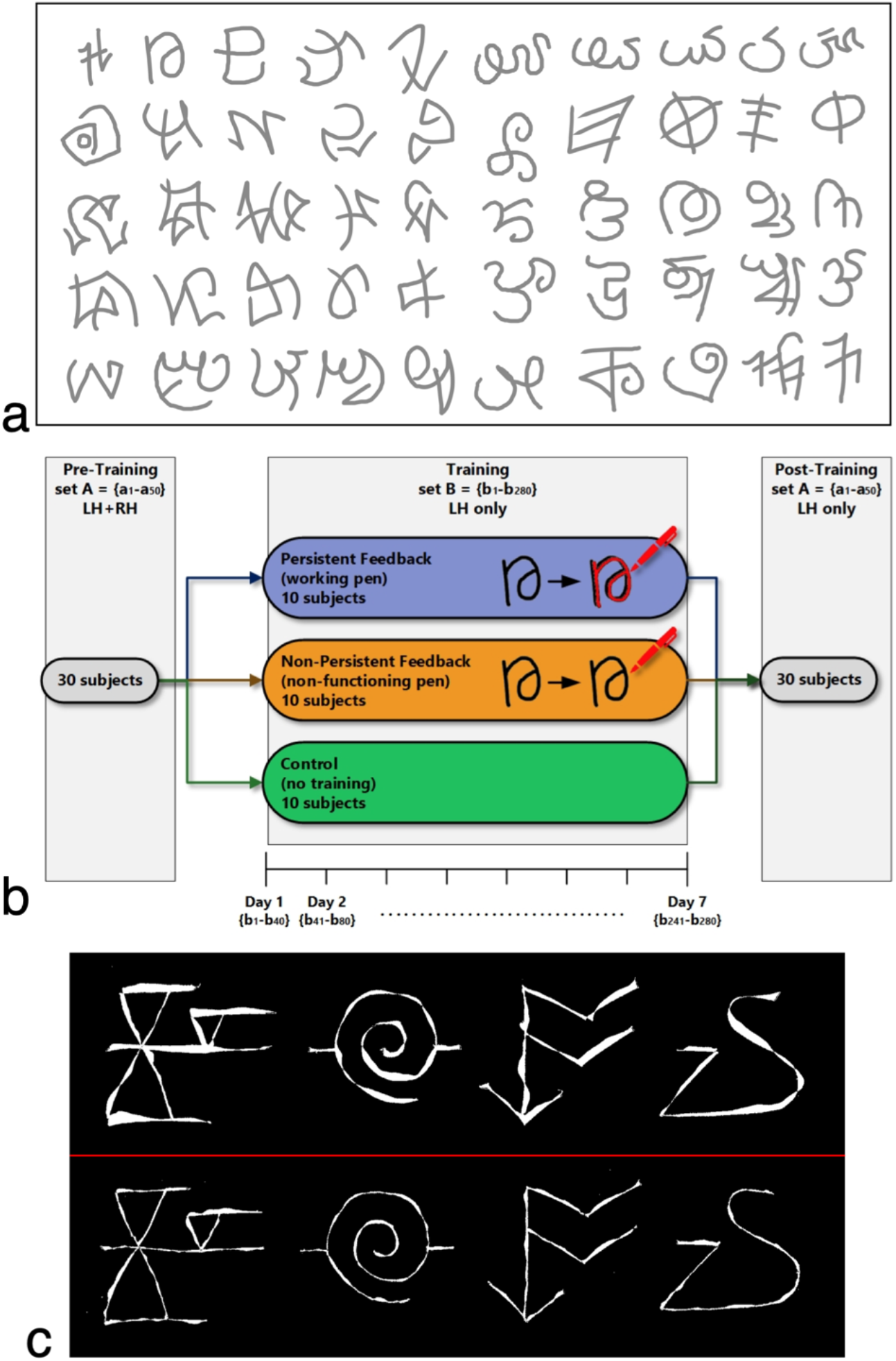
Study methods. (a) Sample patterns subjects were asked to trace in the experiment. (b) 30 subjects participated in a pre-training session, where they were asked to trace patterns in set A with their left and then right hand. The subjects were then randomly split into three training groups. Participants in the Persistent and Non-Persistent Feedback groups were instructed to trace with their non-dominant left hand 40 patterns a day from set B for 7 consecutive days (patterns across training days were non-overlapping). The Control group was asked not to trace during training. All 30 subjects then performed a post-training tracing session over the original patterns from set A using their left hand. (c) The upper panel shows difference in area between four pre-training tracings and the corresponding reference patterns, while the lower panel shows the differences between post-training tracings and the reference shapes, all for the same participant.

### Participants

Thirty college students (14 females; 16 males; mean age across all 30 subjects: 20 years) participated in our study. Each individual was asked about their handedness and any who reported being left-handed, ambidextrous, or frequent users of their left hands for any fine-motor tasks were not included in our study.

### Procedure

Every participant was administered a pre-training test comprising fifty unique letters printed on A4 sheets of paper. They were asked to report any patterns which were familiar to them. With a fine-tipped (0.3 mm) red felt marker, they were asked to trace the images directly on the page, using their left (non-dominant) hand, spending up to 2-3 seconds per image. After tracing with their left hand, participants repeated the test (on a new set sheets), but this time with their right (dominant) hand. Following these two tracing sessions, participants were randomly assigned to three equal-sized groups of ten people each.

#### Group 1: ‘Persistent visual feedback’

Members of group 1 were given a packet comprising seven sets of ten pages each, with each page having four printed patterns. Each of the 280 training patterns were unique and different from the patterns used in the pre- and post-training testing sessions. Participants were told to trace over the patterns of one set per day using only their left-hand and the red felt marker that they were given during the pre-training test. They were told to avoid any extra training or fine visuo-motor tasks with their left hand.

#### Group 2: ‘Non-persistent visual feedback’

Participants in group 2 were given the same packet as those in group 1. The only difference between the two groups was that group 2 received a non-functioning pen to trace the images. This meant that no marks were left on the page as they traced over the patterns. As for group 1, participants in this group were requested to commit to doing the tracings daily (one set per day) across the span of the training period.

#### Group 3: ‘Controls’ (no training)

The control group members were given no training packet and were asked to use their left hand as they normally would, but refrain from undertaking any drawing or other activities involving fine visuo-motor coordination with that hand.

At the end of the training week every participant returned to the laboratory for a post-training testing session. Using their left-hand, they were asked to trace the same fifty patterns that they had traced in the pre-training session. Figure 1b summarizes the overall study protocol.

### Data Analysis

Our analyses were intended to characterize performance along two dimensions: tracing *error* and tracing *steadiness*. Reduction in tracing error reflects the ability to coordinate hand movement in congruence with the reference pattern and is thus an accepted measure of visuo-motor skill improvement (27–31). Steady traces are formed by movement with minimal acceleration change (minimum jerk (33)) and steady exertion of muscle force (34), whereas an increase in acceleration change rate results in less smooth movements with discontinuities which become manifest as unsteadiness in traces. Since smooth movements are thought to be controlled offline (through “global motion planning” (35)), and not affected by the utilization of online visual feedback, steadiness is considered a measure of motor skill.

### Tracing Error

We define two metrics to quantitatively assess subjects’ tracing error:

#### Number of crossovers

Defined as the number of times the drawn curve crosses the reference curve. Two curves that are exactly alike and perfectly superimposed will yield a value of 0 while curves that are imperfect copies will result in multiple intersections, and therefore a high value. This metric has a long history of use in the study of drawing/motor skills (36–38). The number of crossings were counted by one of the authors (ZE), using a blind paradigm (i.e. without knowledge of subject identity, training condition, or time point of produced patterns they were scoring).

#### Area between curves

Defined as the total area of spaces between the reference curve and the tracing (see figure 1c for sample). We digitized all of the tracings and reference figures and superimposed each tracing on the corresponding reference figure. Inter-curve area was computed using a computer program that counted all pixels included in the cross-over spaces between the reference and drawn curves. Two curves that are exactly alike and perfectly superimposed will have no spaces between them, leading to a computed area of zero, while curves that are imperfect copies will result in a higher returned value.

### Tracing Steadiness

Five naive raters (non-overlapping with the participants who had produced the tracings) were presented with pairs of tracings on a computer screen. Each pair consisted of a given subject’s corresponding tracings from pre- and post-training sessions. The left-right positioning of pre- and post-training tracings was randomized across trials. For each pair, raters were asked to score on a five-point scale which tracing looked as if it had been drawn with a steadier hand (1: left side tracing much steadier; 2: left side tracing a little steadier; 3: no difference between the steadiness of the tracings on the two sides; 4 right side tracing a little steadier; 5: right side tracing much steadier). After each rater had scored the entire set, their non-’3’ responses (i.e. those which indicated a difference in steadiness between the two sides) were then compared to an answer key to determine whether or not the side they had selected as steadier was from the pre- or post-training period.

## Results

### Baseline Pre-training Performance

To establish that all three subject groups entered the experiment with similar tracing skills, we compared their baseline tracing performance prior to training. As figure 2a shows, we found no main effect of Group for pre-training tracing performance using the right dominant hand (RDH), as measured by the ‘number of crossings’ error metric (single-factor ANOVA: F = 2.54, df = 2, p = .097). Similarly, the three subject groups had comparable baseline tracing performance when using their left non-dominant hand (LNDH), as measured by both the ‘number of crossings’ (single-factor ANOVA: F = .853, df = 2, p = .437) and ‘area’ (single-factor ANOVA: F = 1.294, df = 2, p = .291) metrics, respectively. Overall, we found no effect of group in pre-training tracing skill, suggesting that our random assignment of participants to the three training groups had not created inadvertent biases for the dominant or non-dominant hand. Not surprisingly, tracing with the RDH far outperformed tracing with the LNDH prior to beginning the training regimen (two-way paired t-test: t_[29]_=10.574, p=1.843E-11).

**Figure 2.**
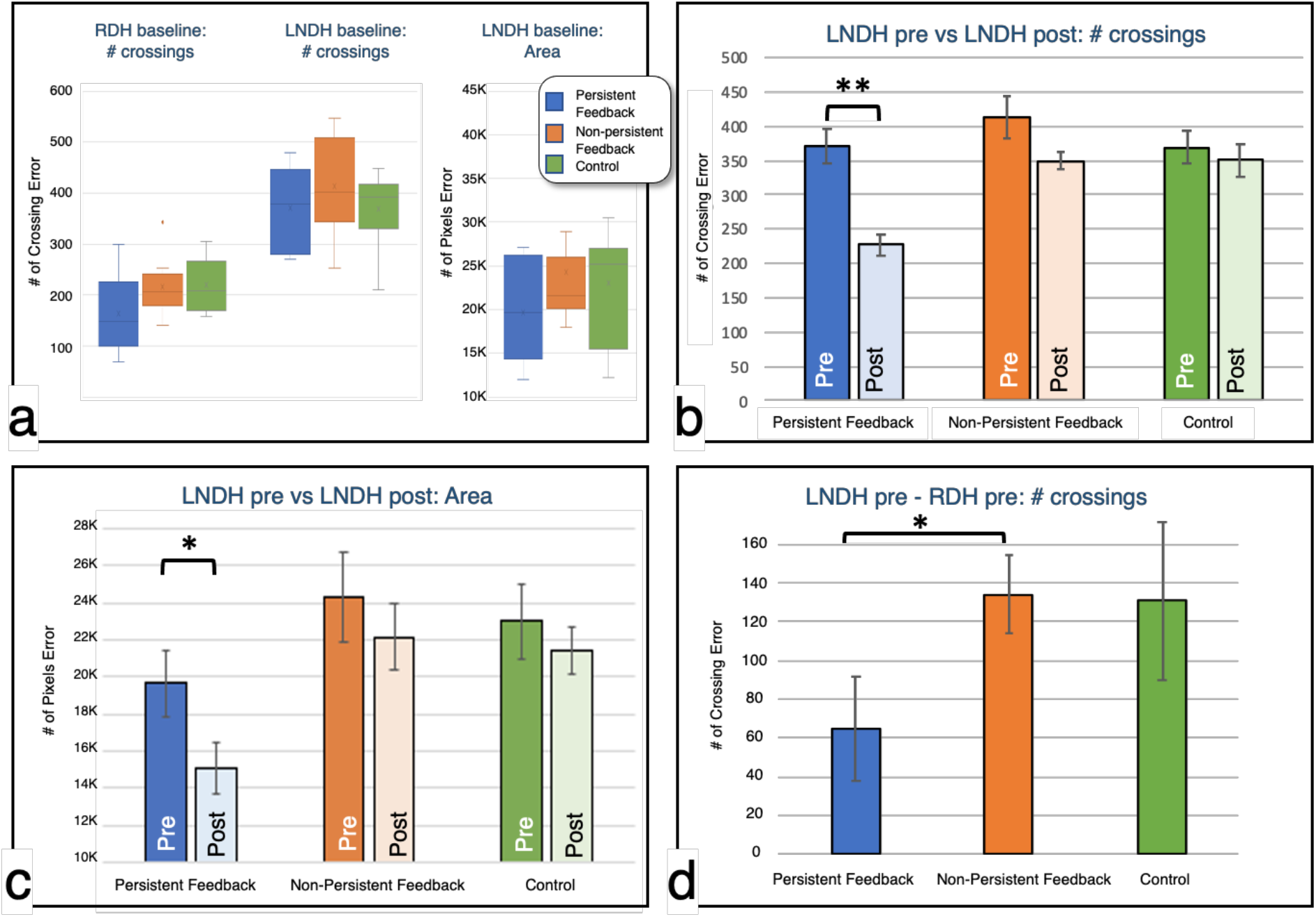
(a) Subjects were assigned to one of the three participant groups randomly: baseline tracing skill, quantified as the ‘number of crossing’ error pre-training, is comparable across subject groups for both the right- and left-hand, indicating no one group was biased with better pre-training skills. (b) and (c) The Persistent Feedback group is the only one that shows statistically significant tracing improvements as a result of their training regimen, for both the number of crossings error and the area metrics. (d) Difference between the LNDH Post-Training error and the RDH Pre-Training error shows that Persistent Feedback training is the only training regimen that leads to LNDH improvements that approach RDH baseline accuracy (i.e. mature tracing performance). Error bars indicate SEM.

### Effect of training-type on tracing error

To assess the effectiveness of our different training programs for improving visuo-motor tracing performance with the LNDH, we compared pre- and post-training error metrics across subject groups. Improvements would be reflected as reduction in error when comparing post- to pre-training patterns. A repeated measures ANOVA on the ‘number of crossings’ data (figure 2b) revealed a significant main effect of training (F = 17.791, df = 1, p < 0.01) and a significant training-by-group Interaction (F = 4.2103; df = 2; p = 0.026), suggesting that training improved performance in some, but not all subject groups. Post-hoc paired comparisons using paired two-tailed t-tests revealed a significant effect of training for the *Persistent Feedback* group (t_[9]_= 4.7925, p= <0.01), but not for the *Non-Persistent Feedback* (t_[9]_= 1.8956, p= 0.091) or *Control* (t_[9]_= .6346, p= 0.542) groups. Consistent with these findings, the two-tailed pairwise comparisons on the ‘area’ data (figure 2c) also revealed a significant effect of training for the *Persistent Feedback* group (t_[9]_= 2.495, p= 0.034), but not the *Non-Persistent Feedback* group (t_[9]_= .625, p= 0.548), or the *Control* group (t_[9]_= .865, p= 0.409).

To assess the extent of improvement that is observed for the left hand, we quantified the error differential between post-training left-hand proficiency and baseline right-hand performance (figure 2d). We found that these differential error-rates were smaller for the *Persistent Feedback* group than for the *Non-persistent* and *Control* groups (Persistent vs Non-persistent: One-tailed t-test: t_[18]_= -1.868, p= .039; Persistent vs Control: One-tailed t-test: t_[18]_= -1.605, p= .063), whereas the non-persistent feedback group did not differ significantly from the control group (One-tailed t-test: t_[18]_= .084, p= .467).

Figure 3 shows the data at a finer granularity; it comprises scatterplots of pre- versus post-training performance for each of our three training regimens, with every data point corresponding to a single subject. For each plot, a data point above, on or below the diagonal corresponds, respectively, to worse, identical or better post-training performance relative to the pre-training value. Given the groups’ comparable baseline performance, the distribution of subjects’ pre-training performance (along the x-axis) is similar across the three subject groups, but the post-training outcomes yield very different scatterplots. All ten participants who received *Persistent Feedback* had a substantial reduction in tracing mistakes, as measured by number of crossings, and all but one improved on the area metric (figures 3a and 3d). In contrast to the downward shift observed for all subjects in the *Persistent Feedback* group, an assessment of the pre- versus post-training errors for subjects in the *Non-persistent Feedback* and *Control* groups exhibits no clear downward shift for both the number of crossings and area metrics. These data show that instantaneous visual and motor feedback on their own (as in the *non-persistent feedback* regimen) are not sufficient for improving tracing accuracy. Such improvement, characterized both by a reduction in the number of crossings and in the area between the reference and produced patterns, requires persistent visual feedback.

**Figure 3.**
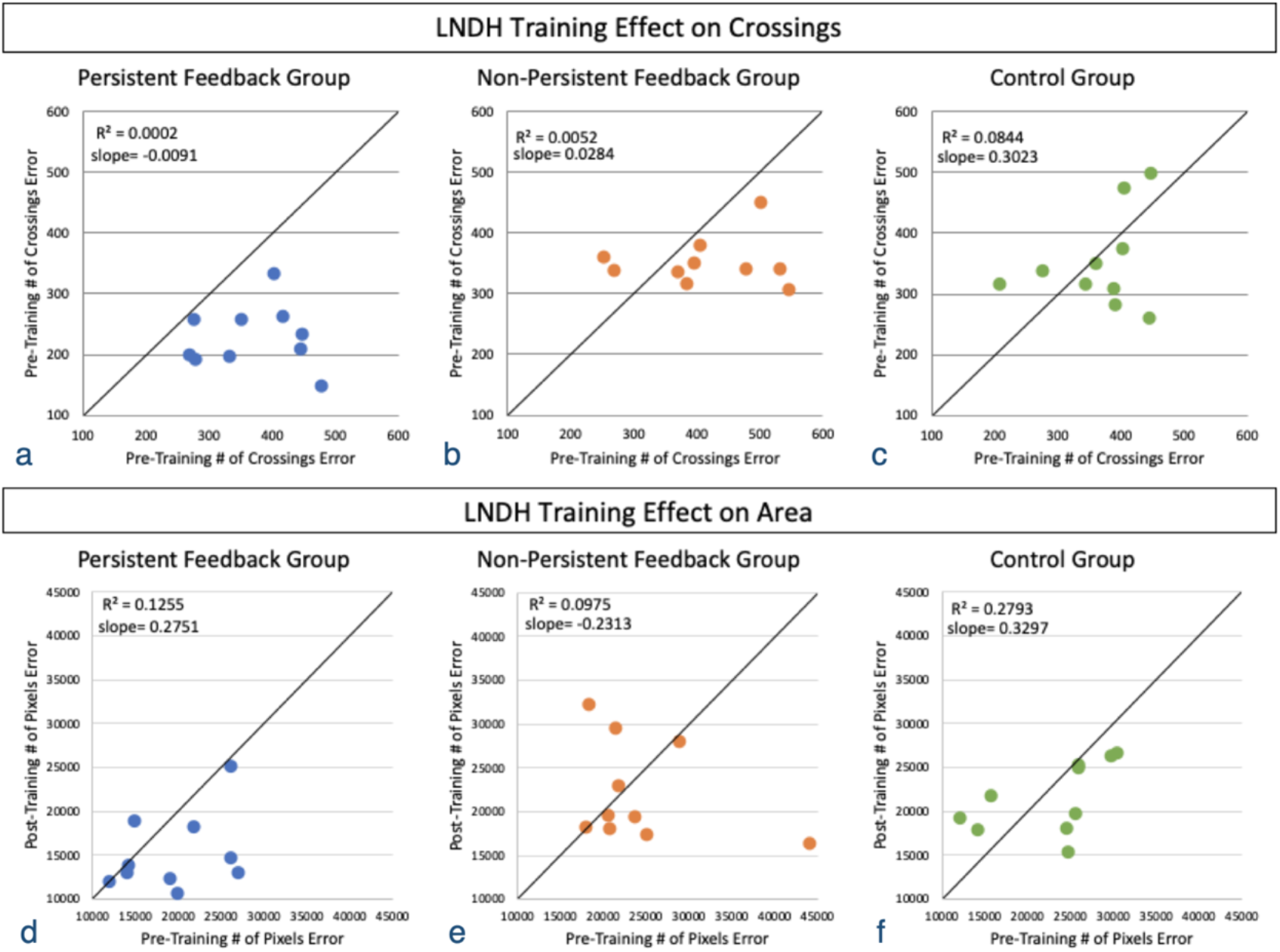
‘Number of crossings’ error (top) and ‘area’ (bottom) scores are plotted to visualize post- (y-axis) versus pre- (x-axis) tracing scores for each subject, represented as individual points. Given that the metrics used represent errors, larger scores correspond to worse tracing performance. Thus, points that lie below, on or above the diagonal indicate subjects with improved, unchanged, and worse post- versus pre-training tracing performance. Virtually every subject in the Persistent Feedback group shows improvement whereas subjects in the Non-persistent feedback and Control groups are more distributed, with more subjects exhibiting worse post-training accuracy than their pre-training accuracy.

### Effect of training-type on steadiness

Figure 4 shows samples of pre- and post-training tracings from each subject group. A cursory inspection of the drawings suggests that there is not necessarily a one-to-one correspondence between a given trace’s accuracy scores and its rated steadiness, i.e. inaccurate lines can be drawn steadily. As the sample tracings in the upper panels of figure 4 suggest, participants in the *Non-persistent* feedback group appear to exhibit improved steadiness from pre- to post-training, despite their error scores not reflecting this. Controls do not show a clear pattern of progress in their left-handed tracing steadiness.

**Figure 4:**
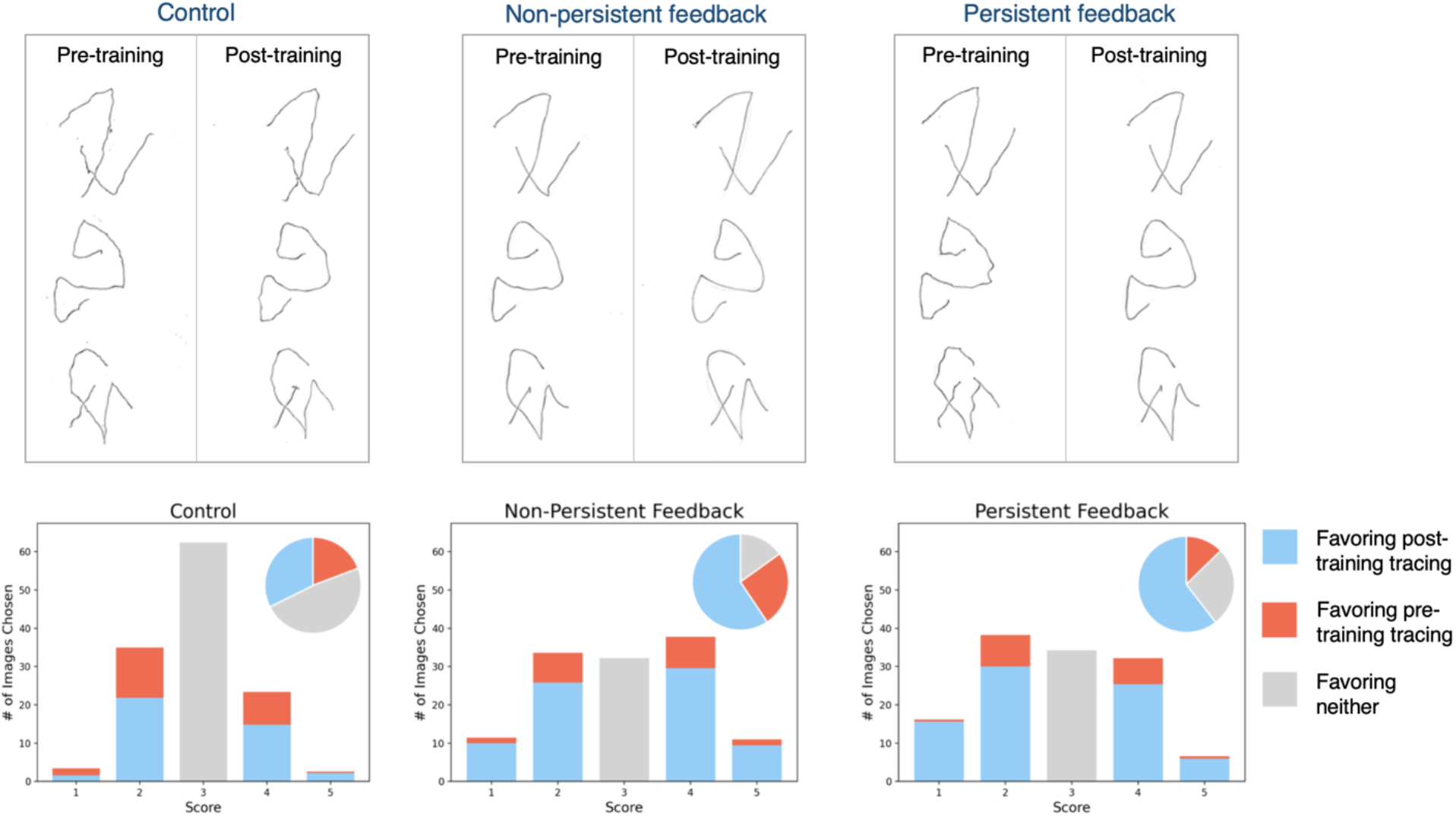
(Upper panels) Sample tracings pairs that ‘steadiness’ raters were shown. Each pair shows one pre- (left) versus post- (right) training tracing of one subject from each group. Visual inspection shows that, in terms of line steadiness, both the Persistent and Non-Persistent training resulted in improvements in tracing steadiness, whereas Control group subjects did not improve. (Lower panels) Average Steadiness scores of five raters. Control group’s tracings were rated mostly as equal steadiness when comparing pre- and post-training tracings, while the other two subject groups had less indeterminate ratings and more post-training tracings identified as steadier.

We enlisted the participation of five individuals naive to the purpose of this study, to serve as raters of tracing steadiness. A summary of their steadiness ratings is shown in the lower panels of figure 4. Tracings of *Control* participants received the largest number of ‘3’ scores suggesting that they had similar steadiness pre- versus post-training period (Binomial tests to determine whether proportion of ‘3’ scores in the persistent and non-persistent conditions were less than those in the control condition yielded p values < 0.05 for all raters individually and as a group). Additionally, post-training tracings from the *Persistent Feedback* as well as *Non-Persistent Feedback* groups were identified as steadier than the pre-training tracings (Binomial tests for each rater show that proportion of post-training tracings assessed to be more steady is significantly higher than the proportion of pre-training tracings, p < 0.05 for all raters). Finally, the data reveal that the proportion of non-3 responses in favor of post-training tracings is not significantly different for the *Persistent* versus *Non-persistent* feedback conditions (Chi-squared test for each rater yields p > 0.05).

## Discussion

Extensive research has examined the interplay between motor skill and visual feedback (7,39–44). The study presented here builds on this rich body of work to investigate what aspects of feedback play a role in improving visuo-motor performance. Specifically, we have examined whether visuo-motor tracing proficiency is impacted differently by persistent versus transient visual feedback. Our data support the following inferences:

### 1. Training with persistent visual feedback, but not transient feedback, leads to an improvement in visuo-motor accuracy (i.e. reduction in error)

This finding is consistent with a hypothesized feedback loop between pattern perception and production, driven by temporally coupled visual and motor information (45–49). The resulting visuo-somato-motor representation facilitates generation of accurate motor programs. The observed improvements generalize to test patterns that are different from those used in training. Also, since our control group showed no significant improvements in skill, we conclude that the results in the persistent feedback group are not merely a consequence of the passage of time, or the experience of taking the pre-assessment test.

### 2. Even short durations of training with persistent feedback are capable of significantly improving visuo-motor accuracy

Our data show that short training sessions (approximately 10 minutes each) across seven days of the *persistent* visual feedback regimen lead to significant improvement in skill. It remains to be seen whether more extended training in the non-persistent feedback condition would eventually result in improvements. We conjecture that, even if extended, this latter regimen is unlikely to match the outcomes of the persistent feedback regimen since recent neuroimaging studies suggest that the two training procedures may recruit different neural pathways (49). While our data reveal the effectiveness of brief training for improving non-dominant hand tracing skills, we do not know how long-lasting these improvements are. Past studies provide some cause for optimism. It has been reported that even modest training of ∼200 minutes can induce substantial improvements in a Precision Drawing Task (50) using the non-dominant hand of healthy adults (51). As the authors suggest, retention of these improvements may be supported by increased connectivity between bilateral sensorimotor hand areas and a left-lateralized parieto-prefrontal praxis network. The longevity of improvements brought about by short training protocols has important implications for rehabilitation of hemiplegic stroke survivors, 20-30% of whom never regain control of their dominant hand for daily tasks requiring a high level of precision (52,53).

### 3. Steadiness of the produced trace is enhanced by training involving either kind of visual feedback - instantaneous as well as persistent

Steadier lines are a characteristic of well-trained motor sequences that correspond to smoother movements (i.e. minimize movement jerk (33,54)). Improved motor skill is associated with more coarticulated movements, such that rather than generating movement units sequentially, each unit is influenced by the anticipated adjacent unit, resulting in their spatial and temporal overlap (35). In fact, the development of the stereotypical smoothness maximization movement specifically depends on observing one’s hand as it moves relative to a target (55), a condition that was fulfilled by both the persistent and non-persistent training regimens, but not by the control group who did not engage in training, and indeed showed no improvements in steadiness.

Taken together, the differential pattern of results that we find between the ‘error’ and ‘steadiness’ metrics can be explained by the former relying on the link between observing the dynamically emergent pattern and the co-occurring motor sequence relative to the reference pattern, while the latter relies only on acquiring a motor sequence planning strategy which results in more articulated movement.

Recent neuroimaging studies in the domain of writing proficiency have suggested that pattern production is a complex visuomotor behavior that involves the concurrent recruitment of occipitotemporal cortex, as well as downstream parietal and motor regions (48). According to this idea, pattern production establishes and strengthens functional pathways between visual and motor areas that are co-activated during motor action, thus facilitating the use of both visual and motor information during pattern perception and production (45). Importantly, such interdependent visuo-motor representation is specific to continuous (i.e. persistent) co-occurrence of visual and motor feedback, as typing does not lead to the same visuo-motor connections that handwriting does (56). Our paradigm builds on these previous studies by focusing on the continuous visual feedback that stems from the produced pattern (49), instead of assuming a-priori that on-line visual feedback for visuo-motor skill improvement relies primarily on information originating from observing one’s hand as it moves relative to a target. Indeed, the consistently differential accuracy improvements that we find between the *persistent visual feedback* versus *non-persistent visual feedback* conditions align well with the aforementioned model; in the non-persistent visual feedback training, subjects only see the static reference pattern throughout the production. The co-occurrence of the dynamically emerging pattern and motor production is thus omitted, and in turn, so is the co-activation and consolidation of those pathways.

Uncovering the neural components underlying mechanisms for acquiring pattern production skills extends our understanding of the interplay between vision and motor actions for rich representations, and brings with it many real-world applications, from development of interactive educational technology, to revising assessment and rehabilitation programs for stroke patients, and even for the development of biomimetic multimodal algorithms for motor planning and execution of tasks in robotic systems. Of particular interest to us is the field of rehabilitation of children with visual impairments, particularly those with atypical visual development. Our research group has had the opportunity to conduct a unique program in India in which we provide surgical intervention for children with treatable congenital blindness, and then study how their visual system learns to make sense of the world when they begin to see late in life (www.projectprakash.org). Despite the effective reversal of their blindness, these children, and many like them worldwide, struggle with significantly impaired visuo-motor skills (figure 5), compromising their ability to undertake normal schooling. An effective method for improving these skills will be of profound importance for such children. Our findings on the role of persistent visual feedback for enhancing motor accuracy, taken in the context of other recent studies about the role of dynamically emergent and co-occurring visual and motor feedback for pattern production and recognition, suggest that even such late-sighted children may be able to learn critical writing skills and engage in drawing tasks, if that learning emphasizes, rather than omits, the reliance on temporally emergent and persistent visual feedback.

**Figure 5.**
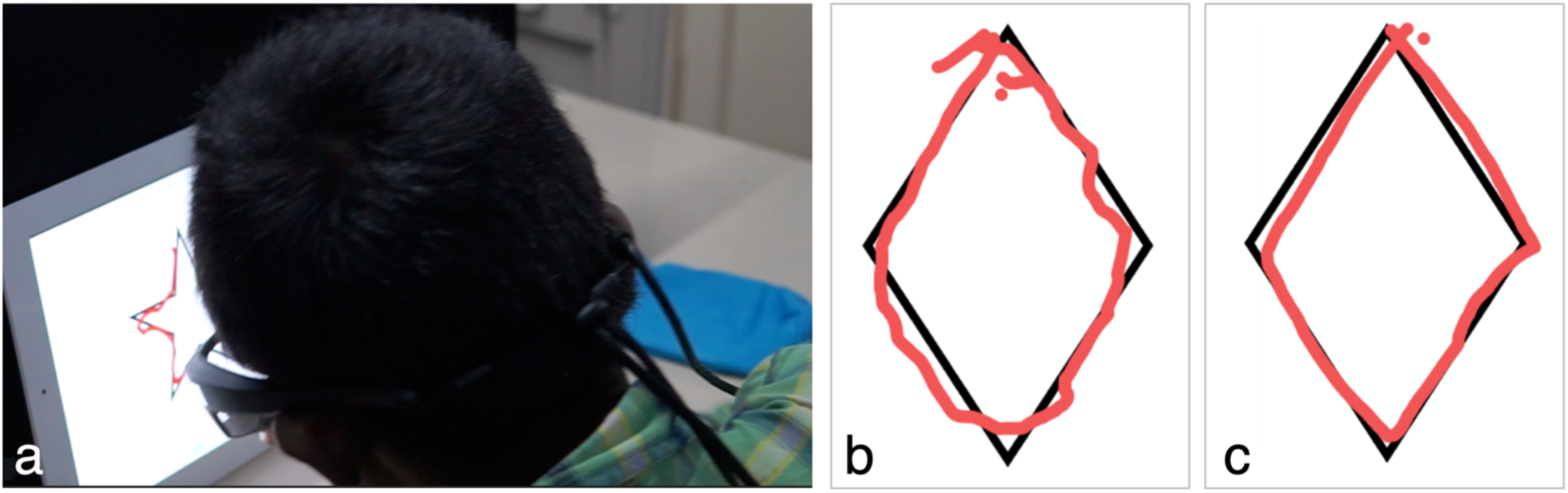
(a) A 12yo Prakash child performs a tracing task after treatment for bilateral congenital cataracts. (b) A sample tracing of a diamond by this child one-month post-treatment (pre-operative bilateral acuity was 20/3200, acuity at time of tracing experiment was 20/600). (c) Tracing by a control 10yo male with typically developing vision while wearing blur goggles that impose a 20/500 bilateral acuity.

Overall, our results suggest that the continual act of moving the hand through various patterns while leaving a temporally persistent trace, and observing the congruence of the resulting pattern trajectory with the reference pattern can improve proficiency on a visuo-motor coordination task. This is a step towards identifying the types, and the mechanistic underpinnings, of visual cues that are most effective for visuo-motor training.

## Notes

### Competing Interest Statement

The authors have declared no competing interest.

